# Non-canonical localization of RubisCO under high light conditions in the toxic cyanobacterium *Microcystis aeruginosa* PCC7806

**DOI:** 10.1101/695841

**Authors:** Tino Barchewitz, Arthur Guljamow, Sven Meissner, Stefan Timm, Manja Henneberg, Otto Baumann, Martin Hagemann, Elke Dittmann

**Affiliations:** Department of Microbiology, Institute for Biochemistry and Biology, University of Potsdam, Karl-Liebknecht-Str. 24/25, 14476 Potsdam-Golm, Germany; Department of Plant Physiology, Institute Biological Sciences, University of Rostock, Albert-Einstein-Str. 3, 18059 Rostock, Germany; Department of Zoophysiology, Institute for Biochemistry and Biology, University of Potsdam, Karl-Liebknecht-Str. 24/25, 14476 Potsdam-Golm, Germany

**Author notes:** Correspondence should be addressed to Prof. Dr. Elke Dittmann, University of Potsdam, Institute of Biochemistry and Biology, Department of Microbiology, Karl-Liebknecht-Str. 24/25, 14476 Potsdam-Golm Germany.

**Keywords:** cyanobacteria, RubisCO, microcystin, inorganic carbon fixation

## Abstract

The frequent production of the hepatotoxin microcystin and its impact on the life-style of bloom-forming cyanobacteria are poorly understood. Here we report that microcystin interferes with the assembly and the subcellular localization of RubisCO, in *Microcystis aeruginosa* PCC7806. Immunofluorescence, electron microscopic and cellular fractionation studies revealed a pronounced heterogeneity in the subcellular localization of RubisCO. At high cell density, RubisCO particles are largely separate from carboxysomes in *M. aeruginosa* and relocate to the cytoplasmic membrane under high-light conditions. We hypothesize that the binding of microcystin to RubisCO promotes its membrane association and enables an extreme versatility of the enzyme. Steady-state levels of the RubisCO CO_2_ fixation product 3-phosphoglycerate are significantly higher in the microcystin-producing wild type. We also detected noticeable amounts of the RubisCO oxygenase reaction product secreted into the medium that may support the mutual interaction of *M. aeruginosa* with its heterotrophic microbial community.

## Introduction

Bloom-forming cyanobacteria are infamous for the production of toxins and constitute a serious threat for humans and animals. Among the cyanobacterial toxins, microcystin (MC) stands out as the most widely encountered type of toxin (Dittmann, Fewer, & Neilan, 2013; Huisman et al., 2018). The toxicity of the cyclic heptapeptide was attributed to the inhibition of eukaryotic-type protein phosphatases of the families PP1 and PP2A (Goldberg et al., 1995). While exposure to MC is often fatal for animals, a primary role of MC as feeding deterrent is increasingly under debate (Rohrlack, Dittmann, Borner, & Christoffersen, 2001). A phylogenetic analysis of MC biosynthesis genes from distant cyanobacterial genera revealed that the genes were already present in the last common ancestor of all recent cyanobacteria and hence prior to the evolution of eukaryotes (Rantala et al., 2004). Moreover, MCs are primarily intracellular toxins and the levels dissolved in water typically do not exceed critical levels of toxicity (Dittmann et al., 2013).

Although MCs were incidentally detected in terrestrial ecosystems (Kaasalainen et al., 2012), there is a striking prevalence of these toxins in bloom-forming freshwater cyanobacteria such as *Microcystis, Planktothrix* and *Dolichospermum* (Dittmann et al., 2013; Huisman et al., 2018). These genera belong to diverse subclades of the cyanobacterial phylum, yet they share a number of common features. Species showing mass developments in lakes often have the ability to migrate vertically in the water column thereby experiencing fluctuating light conditions (Rabouille, Thebault, & Salencon, 2003). This buoyancy regulation is based on the presence of gas vesicles and the formation of multicellular colonies or filaments. The phototrophic aggregates are extensively colonized by specific communities of heterotrophic bacteria (Woodhouse, Ziegler, Grossart, & Neilan, 2018), which at least in part have enhancing effects on the growth of the cyanobacteria (Berg et al., 2009). In the upper layer of the water column where they are exposed to high irradiances, photosynthetically active cyanobacterial populations generate high levels of O_2_ and a high pH while inorganic carbon (Ci) becomes scarce (Havens, 2008); (Sandrini et al., 2016). Bloom-forming species can cope surprisingly well under these self-inflicted extreme conditions considering the fact that CO_2_ fixation and growth of cyanobacteria largely depend on the bifunctional enzyme RubisCO for which CO_2_ and O_2_ represent competing substrates. Among cyanobacteria, robust carbon fixation by RubisCO under limiting C_i_ conditions is ensured by the carbon concentrating mechanism (CCM), whereby cyanobacteria raise the CO_2_ level in the vicinity of RubisCO by a complement of bicarbonate and CO_2_ uptake systems and by encapsulating RubisCO and carbonic anhydrase in carboxysomes (Burnap, Hagemann, & Kaplan, 2015). In this context it is interesting to note that strains of *Microcystis*, in spite of having highly similar core genomes, selectively lost or acquired individual bicarbonate transporters (Sandrini, Matthijs, Verspagen, Muyzer, & Huisman, 2014).

Apart from the buoyancy regulation, another unifying feature of bloom-forming cyanobacterial species is their ability to produce a specific set of specialized molecules. Besides MC, hallmark peptide families in bloom-forming cyanobacteria include cyanopeptolin, aeruginosin, microginin, anabaenopeptin, and aeruginoguanidine (Pancrace et al., 2018; Welker & von Dohren, 2006). In contrast to secondary metabolites in other microorganisms, the peptides are produced from the beginning of the logarithmic growth and are primarily located inside the cells (Long, Jones, & Orr, 2001; Rapala, Sivonen, Lyra, & Niemela, 1997). Each of the peptides is formed by a giant non-ribosomal peptide synthetase assembly line, an individual bloom-forming strain thus encodes several of these multienzyme complexes in parallel and devotes a large part of its resources to the production of these specialized compounds (Welker & von Dohren, 2006).

There is rising evidence for a relationship between MC, RubisCO and the CCM in the bloom-forming species *Microcystis*. Two independent studies have revealed that the MC-producing strain *M. aeruginosa* PCC7806 can grow better under C_i_ limitation than the nontoxic Δ*mcyB* mutant which *vice versa* outcompetes the wild-type strain at high CO_2_ levels (Jahnichen, Ihle, Petzoldt, & Benndorf, 2007; Van de Waal et al., 2011). Moreover, the toxin was shown to bind to a number of proteins in *Microcystis*, among which the large (RbcL) and small (RbcS) subunits of RubisCO and a striking number of Calvin-Benson cycle enzymes such as phosphoribulokinase, phosphoglycerate kinase, fructose-bisphosphate aldolase, and glyceraldehyde-3 phosphate dehydrogenase were identified as predominant binding partners (Wei, Hu, Song, & Gan, 2016; Zilliges et al., 2011). MC-bound RubisCO, in turn, is significantly more stable against protease degradation (Zilliges et al., 2011). Furthermore, a metabolomic comparison of wild-type and Δ*mcyB* mutant extracts revealed major differences in the accumulation of glycolate as a product of the RubisCO oxygenase reaction under high-light conditions (Meissner, Steinhauser, & Dittmann, 2015). This phenotype resembles the observed accumulation of oxygenase products of Rubisco in carboxysome-defect mutants of the model strain *Synechocystis* sp. PCC 6803 (Hackenberg et al., 2012). Metabolomic differences between *M. aeruginosa* PCC7806 and its MC-deficient mutant were generally more pronounced under high-light conditions, the same conditions that stimulate both the expression of MC biosynthesis genes and MC-binding to proteins (Kaebernick, Neilan, Borner, & Dittmann, 2000). Binding of MC to proteins was also observed in field colonies of *Microcystis*, while in the laboratory it is only observed at higher cell densities (Meissner, Fastner, & Dittmann, 2013; Wei et al., 2016).

Here, we have assessed whether differences in the subcellular localization of RubisCO and more specifically a localization inside or outside of carboxysomes may contribute to the pronounced metabolomic differences between *M. aeruginosa* PCC7806 and the Δ*mcyB* mutant under high-light conditions. Our data suggest that localization and assembly of RubisCO in *Microcystis* can deviate from the current textbook view on RubisCO and the CCM among cyanobacteria and provide evidence that MC is interfering with the assembly of the RubisCO complex.

## Results

### Dynamic subcellular localization of RubisCO in Microcystis

To evaluate whether the light-dependent dynamics in the accumulation of RubisCO products in *M. aeruginosa* (Meissner et al., 2015) is due to changes in the subcellular localization of the enzyme, we performed light shift experiments for up to 4 h with cells of the MC-producing wild-type strain (WT) and the MC-deficient Δ*mcyB* mutant, which were pre-grown under low light conditions at ambient air. Steady state levels of the immediate products of RubisCO, 3-phosphoglycerate (3-PGA) and 2-phosphoglycolate (2-PG) were determined using LC-MS. Levels of 3-PGA were around ten-fold higher in the WT strain compared to the Δ*mcyB* mutant, both under low-light and high-light conditions (Fig. 1A). 2-PG, on the other hand, was continuously accumulating independent of the light conditions. Considerable amounts of 2-PG were detected in the culture supernatant and were even exceeding intracellular levels of the metabolite in the Δ*mcyB* mutant (Fig. 1B). Yet, the overall amount of the metabolite was rather low and differences between WT and mutant strain were neglectable. Collectively, these data indicate that the ratio between carboxylation and oxygenation activity of RubisCO differs between the two strains, whereby the RubisCO carboxylase activity is favoured in the MC-producing WT.

**Fig. 1.**
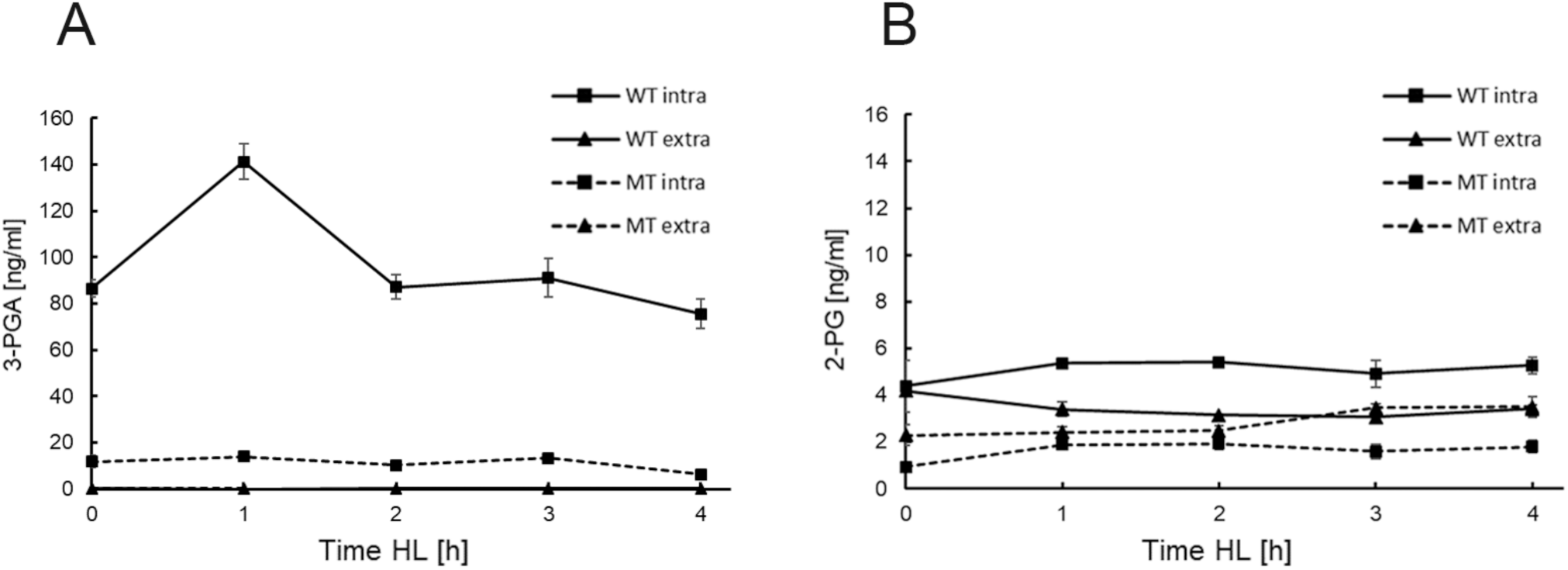
Accumulation of RubisCO products *in vivo*. Liquid chromatography-mass spectrometry (LC-MS) analysis of steady-state levels of 3-phosphoglycerate (3-PGA) and 2-phosphoglycolate (2-PG). Low-light adapted cultures (15 µmol photons m^-2^s^-1^) of *M. aeruginosa* wild type (WT) and the Δ*mcyB* mutant (MT) were exposed to high-light (250 µmol photons m^-2^s^-1^) for up to 4 h. A) Intra- and extracellular steady-state levels of 3-PGA, B) Intra- and extracellular steady-state-levels of 2-PG. Shown are mean values of three biological replicates.

To assess whether the differences in the accumulation of 3-PGA in the MC-producing WT and the Δ*mcyB* mutant are related to differences in the subcellular localization of RubisCO, cellular protein extracts from both strains were separated into soluble and membrane fractions using samples along the 4 h low-light to high-light time course (Fig. 2A). The quality of the separations was verified using an antibody against the key protein of cell division, FtsZ, that yielded signals in the soluble fraction, only (Fig. 2B). Unexpectedly, RbcL was increasingly detected in the membrane fraction under high-light conditions leading to an almost exclusive RbcL localization in the membrane fraction after 4 h high-light conditions (Fig. 2B). The membrane-associated RbcL fraction could be mechanically detached by sonication indicating that RbcL is only loosely associated with membranes in *Microcystis.* Correspondingly, RubisCO activity was only measured in the fraction loosely associated with membranes and not in the membrane fraction with tightly bound proteins after detachment of RubisCO (Fig. S1). The relocation of RbcL from the cytoplasm towards the membrane was more pronounced in the MC-producing WT than in the Δ*mcyB* mutant, in which RbcL notably disappeared after 4 h high-light treatment (Fig. 2 C). The same protein fractions were also tested with an antibody against the major carboxysomal shell protein, CcmK (see methods section for the generation of polyclonal CcmK antibody). This protein was primarily detected in the cytosolic fraction as expected but increasingly appeared in the membrane-associated fraction with enduring high-light treatment (Fig. 2B). Compared to the subcellular localization shift of RbcL, the CcmK relocation to the membrane fraction was lagging behind leading to the non-expected observation that a substantial amount of RbcL and CcmK was found in separate fractions after 1 and 2 h high-light treatment. These data clearly indicate a predominant non-carboxysomal localization of RbcL conditions. The fractions were additionally tested with a specific antibody against RbcS (see methods section for the generation of the polyclonal RbcS antibody). Remarkably, subcellular localization of RbcS was not mirroring the localization of its counterpart RbcL. Instead, RbcS was equally present in the cytosolic and the membrane-associated fraction during the entire light-shift experiment indicating that the two subunits of the canonical hexadecameric RbcL_8_RbcS_8_ RubisCO complex, at least partly, separate from each other in high-light treated cells of *Microcystis*. RbcS also seemed to be more stable than RbcL in the mutant after 4 h high-light treatment suggesting that not only the localization but also the turnover of both proteins differs in *Microcystis*.

**Fig. 2.**
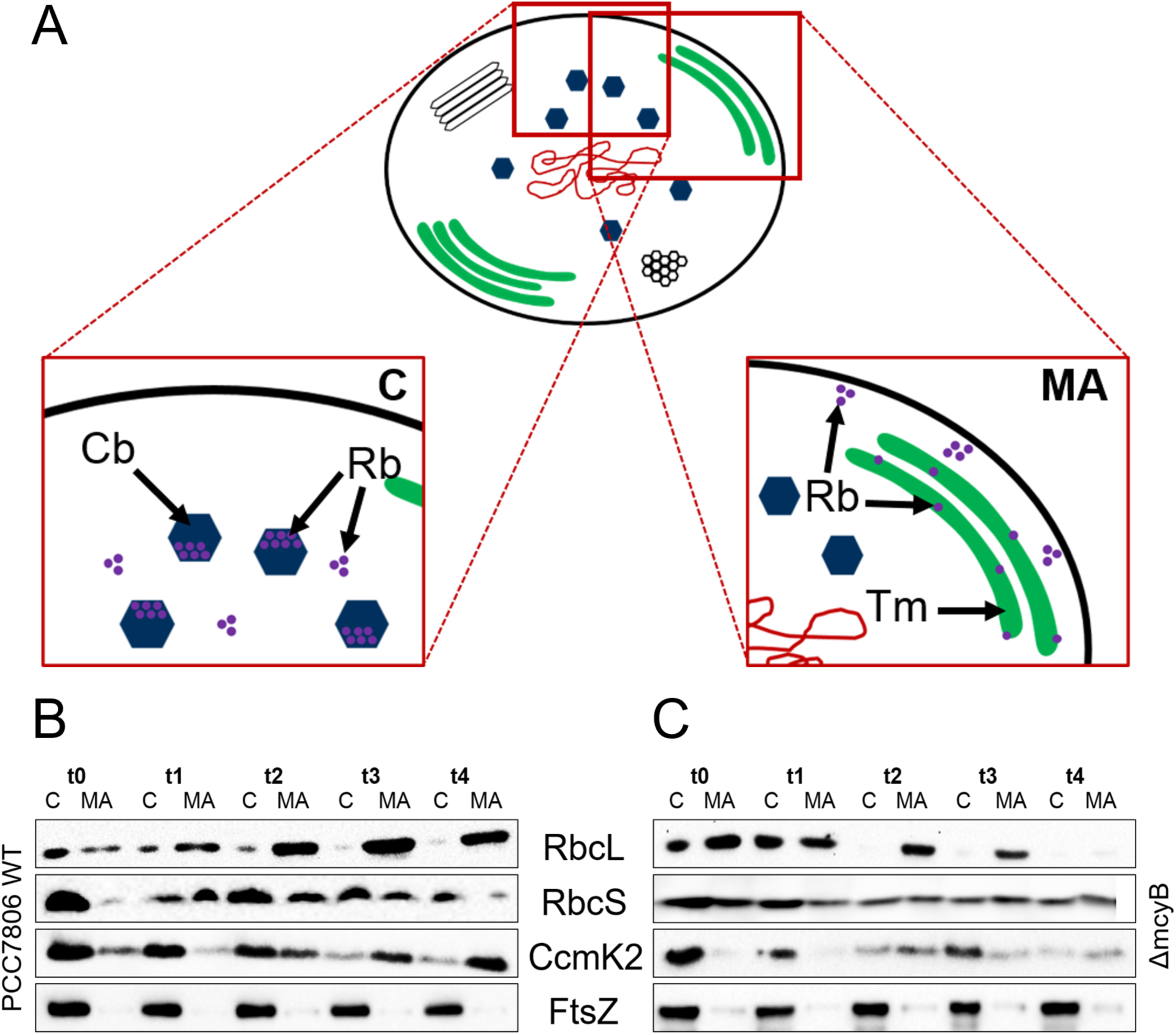
Dynamics of key proteins of carbon fixation during a light shift experiment in *M. aeruginosa* PCC7806. Low-light adapted cultures of *M. aeruginosa* wild type (WT) and the Δ*mcyB* mutant (MT) were exposed to high-light for up to 4 h. Light conditions were as for Fig. 1. (**A)** Schematic representation of subcellular localization of RubisCO in *M. aeruginosa*. RubisCO (Rb) can be localized in the cytosol (C) and encapsulated in carboxysomes (Cb) or associated with membranes such as the thylakoid membrane (Tm) or the cytosolic membrane (MA). **(B-C)** Western blots showing the relocation of RbcL and CcmK from the cytosolic fraction (C) towards the membrane-associated fraction (MA) during 4 h of high-light treatment in the WT and the MT, respectively. RbcS is located both in the cytosolic (C) and the membrane-associated fraction (MA) independent of the light condition. FtsZ serves as cytosolic marker and confirms the separation of cytosolic and membrane-associated proteins.

### RbcL is localized underneath the cytoplasmic membrane in *Microcystis*

To verify the relocalization of RbcL in high-light treated cells by an independent method, we next utilized immunofluorescence microscopy (IFM) to visualize the subcellular localization of RbcL and CcmK in cells along the same high-light time-course. Using an antibody against the carboxysomal protein CcmK as a proxy, the IFM methodology in *Microcystis* was optimized to assure the permeability of carboxysomal structures without completely destroying their integrity. CcmK was primarily detected in the central cytosolic area and became visible as rings in optical sections indicating that CcmK indeed assembles into characteristic carboxysomal structures in *Microcystis*. CcmK signals were occasionally also observed at the periphery of cells (Fig. 3). No specific signals were obtained in control experiments without the primary CcmK antibody (Fig. S2). While the ring-like structure confirmed the integrity of carboxysomes, the diffuse signals support a sufficient degree of permeabilization to reliably detect RubisCO within or leaking out of carboxysomes. We neither observed major variations in the subcellular localization of CcmK during the low-light to high-light shift experiment, nor considerable differences between the MC-producing WT and the MC-free mutant strain (Fig. 3). In contrast, the distribution of RbcL showed a considerable heterogeneity among cells and a remarkable dynamics between low-light and high-light conditions. RbcL was predominantly detected in small spots distributed across the cell in the *Microcystis* WT. The localization of RbcL in larger bodies, indicative of a possible carboxysomal localization, was only observed within a small fraction of cells under low light conditions (Fig. 3A). Remarkably, the small RbcL spots relocated toward the cytoplasmatic membrane with continuing high-light treatment (Fig. 3A). Cytoplasmatic or carboxysomal RbcL signals were virtually absent in these high-light treated cells. In MC-deficient mutant cells, RbcL was more evenly distributed within small distinct spots primarily occurring in the cytosolic area (Fig. 3B). Although a cytoplasmic membrane localization was occasionally observed after 4 h of high-light treatment, the relocation phenomenon was clearly less pronounced in mutant than in WT cells. Again, an apparent carboxysomal localization of RubisCO was only occasionally observed in a small subfraction of cells (Fig. 3B). These data further strengthen the observation that carboxysomes and RbcL are separated in high-light treated *Microcystis*, in particular in the MC-producing WT. While a partial association of RubisCO with the thylakoid membrane has been reported before for the model cyanobacterium *Synechocystis* sp. PCC6803 (Agarwal, Ortleb, Sainis, & Melzer, 2009), localization of RubisCO underneath the cytoplasmic membrane is unprecedented and has never been reported for cyanobacteria.

**Fig. 3.**
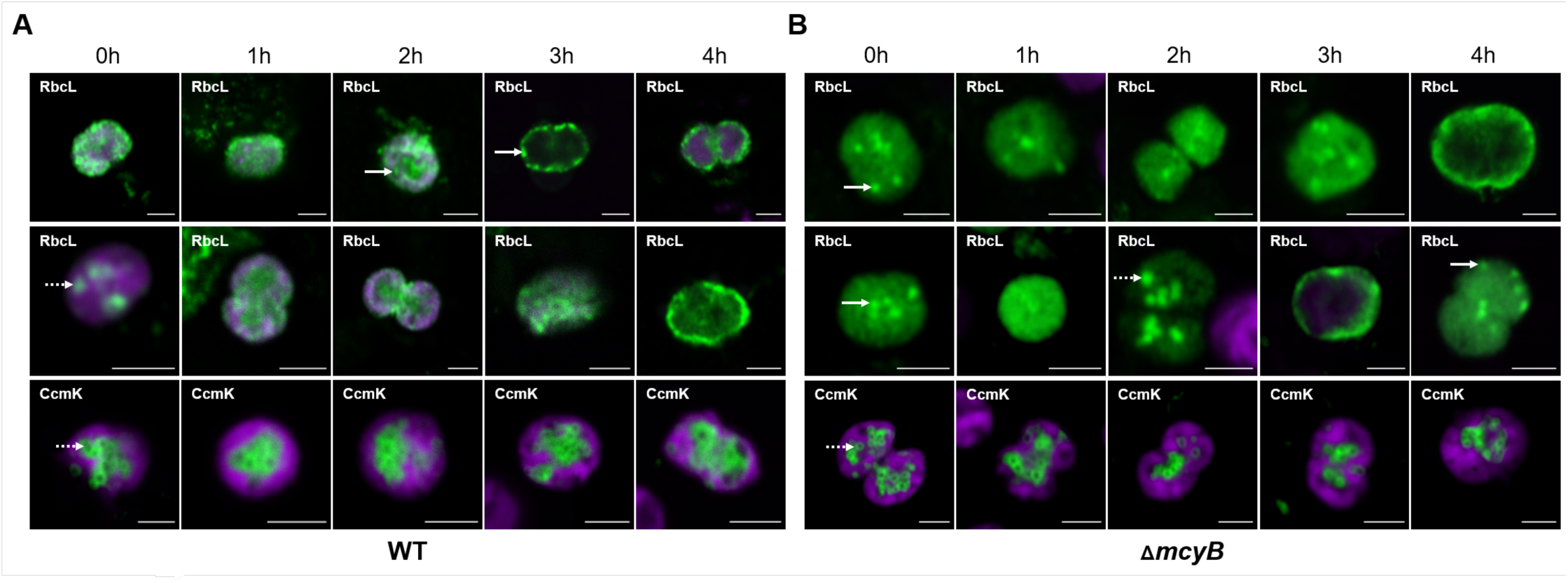
Immunofluorescence micrographs (IFM) visualizing the subcellular localization of RbcL and CcmK in *M. aeruginosa* wild type (WT) and Δ*mcyB* mutant during a light shift experiment. Light conditions were as described for Fig. 1 **(A)** Immunostaining of RbcL and CcmK in *M. aeruginosa* WT cells with respective antibodies. **(B)** Immunostaining of RbcL and CcmK in *M. aeruginosa* Δ*mcyB* mutant cells with respective antibodies. The green fluorescence signal indicates the subcellular localization of RbcL and CcmK, respectively; the purple signal reflects the autofluorescence of thylakoid-associated phycobiliproteins. In the WT, RbcL appears as spots in the cytosol or underneath the cytoplasmic membrane (arrow) or in apparent carboxysomal bodies (dashed arrow), while it shows a homogenous distribution with a few concentrated spots in the Δ*mcyB* mutant. CcmK signals show the characteristic carboxysome structures (arrow) inside the cytosol and some undefined structures at the cytosolic membrane (dashed arrow) in both WT and in the Δ*mcyB* mutant strains. The scale bar in all images is 2 µm.

In order to assess how far the distinct subcellular localization of RubisCO in the MC-producing WT and the mutant relate to the binding of MC to RubisCO, we performed diurnal cultivation experiments (16 h light of 60 µmol photons m^-2^ s^-1^, 8 h darkness) with the WT strain at high cell density (OD_750_: 0.6) and low cell density (OD_750_: 0.3),. As seen in a previous study, MC-binding to proteins was only detected in the high cell density cultures but not in the low cell density cultures (Wei et al., 2016) (Fig. 4B and D). Double-staining of high cell-density cells combining antibodies against RbcL and CcmK revealed a distinct localization of the two proteins at the cytoplasmic membrane and in carboxysomal bodies, respectively, during the day. Even when carboxysomal structures were completely disrupted after formaldehyde fixation, no major portion of RbcL was detected in the cytosol, thereby excluding the possibility that the lack of RbcL signals is due to the insufficient penetration of the RbcL antibody through shell structures (Fig. 4 A). In contrast to high cell density cultures, RbcL and CcmK showed a predominant cytoplasmatic localization at low cell density including a colocalization with CcmK signals, suggesting that RbcL is more frequently located in carboxysomes under these conditions (Fig. 4C). Moreover, only at high cell densities subcellular localization of RbcL differed between day and night samples (Fig. 4). Samples harvested during the dark phase revealed a relocation of RbcL towards the thylakoid membrane facing the innermost cytoplasmic space (Fig. 4A).

**Fig. 4.**
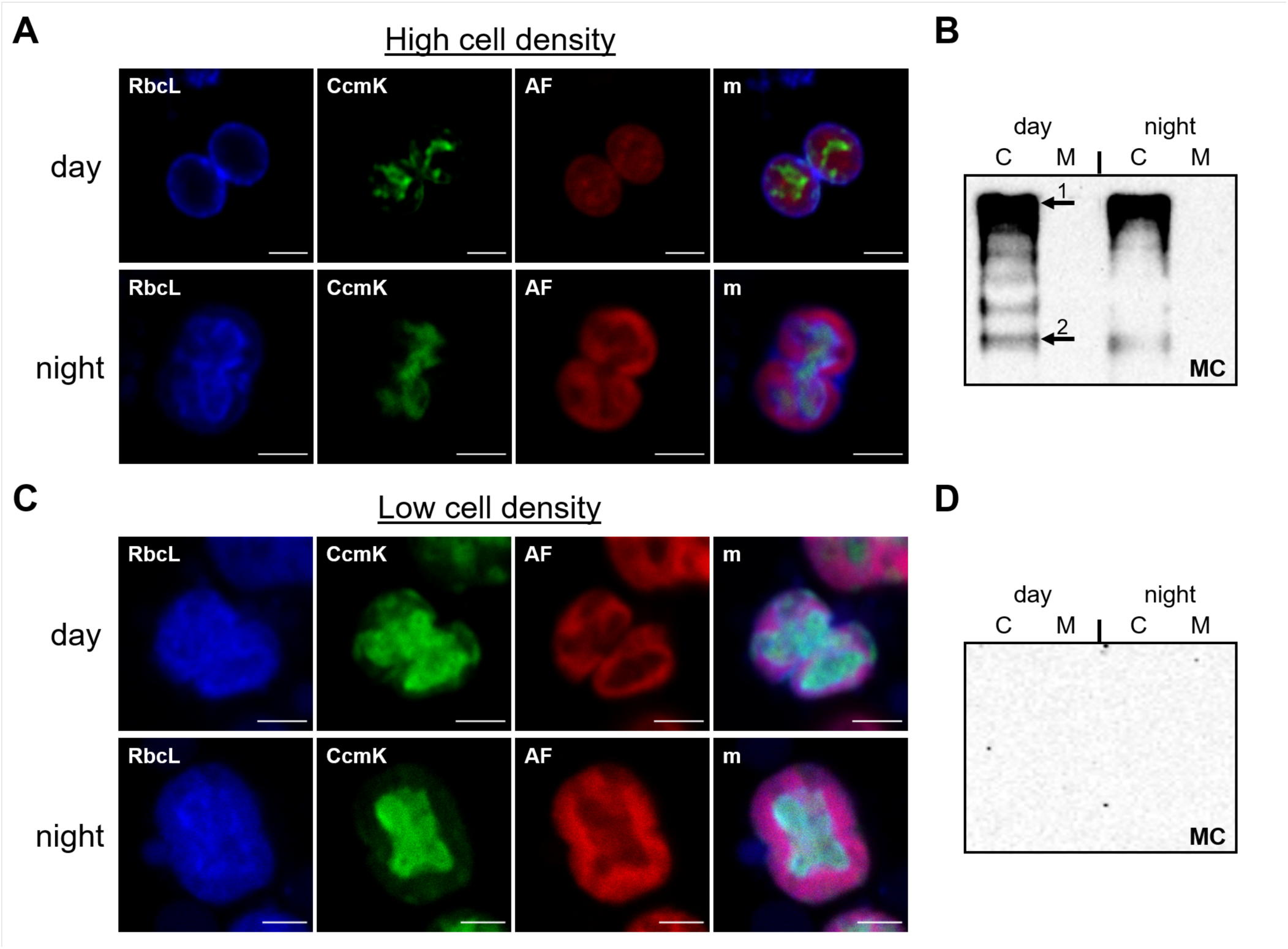
Co-Localization of RbcL and CcmK in *M. aeruginosa* wild type PCC7806 at different cell densities visualized by IFM. **(A and C)** Co-Hybridization with RbcL and CcmK antibodies at **(A)** high cell density (OD_750_: 0.6) and **(C)** low cell density (OD750: 0.3). RbcL is visible in the blue fluorescence channel and CcmK is visible in the green fluorescence channel. AF: red phycobilisome autofluorescence, m: merged image from the 3 fluorescence channels. AF=phycobilisome auto fluorescence, m=merged image from the 3 fluorescence channels. The scale bar in all images is 2 µm. **(B and D)** Western blot detection of protein-bound MC in soluble and membrane protein extracts of **(B)** high cell density cultures and **D)** low cell density cultures, respectively. The strongest MC signals were obtained for an SDS-stable high-molecular mass complex (1) (see Fig. 8 for further analysis), but signals were also observed at the level of RbcL at 52 kDa (2).

We further tested co-localization of RbcL and RbcS in high-light treated WT and mutant cells at high cell density (Fig. 5). The two subunits of RubisCO showed partly distinct localization patterns in the wild type with RbcS predominantly located in the central cytosolic space and near the thylakoids, and RbcL predominantly located underneath the cytoplasmic membrane (Fig. 5A). The presence of both RbcL and RbcS in the membrane-associated fraction (Fig. 2) thus does not necessarily mean that both subunits are found in hexadecameric complexes located at the same membrane. It should be noted, however, that the RbcS antibody was raised against the recombinant monomeric protein, raising the possibility that the antibody does not recognize RbcS well in the hexadecameric context *in situ*. RbcL and RbcS localizations were partially overlapping in the MC-deficient mutant, yet their distribution was still different with RbcL rather evenly distributed and RbcS confined to distinct spots resembling carboxysome structures (Fig. 5).

**Fig. 5.**
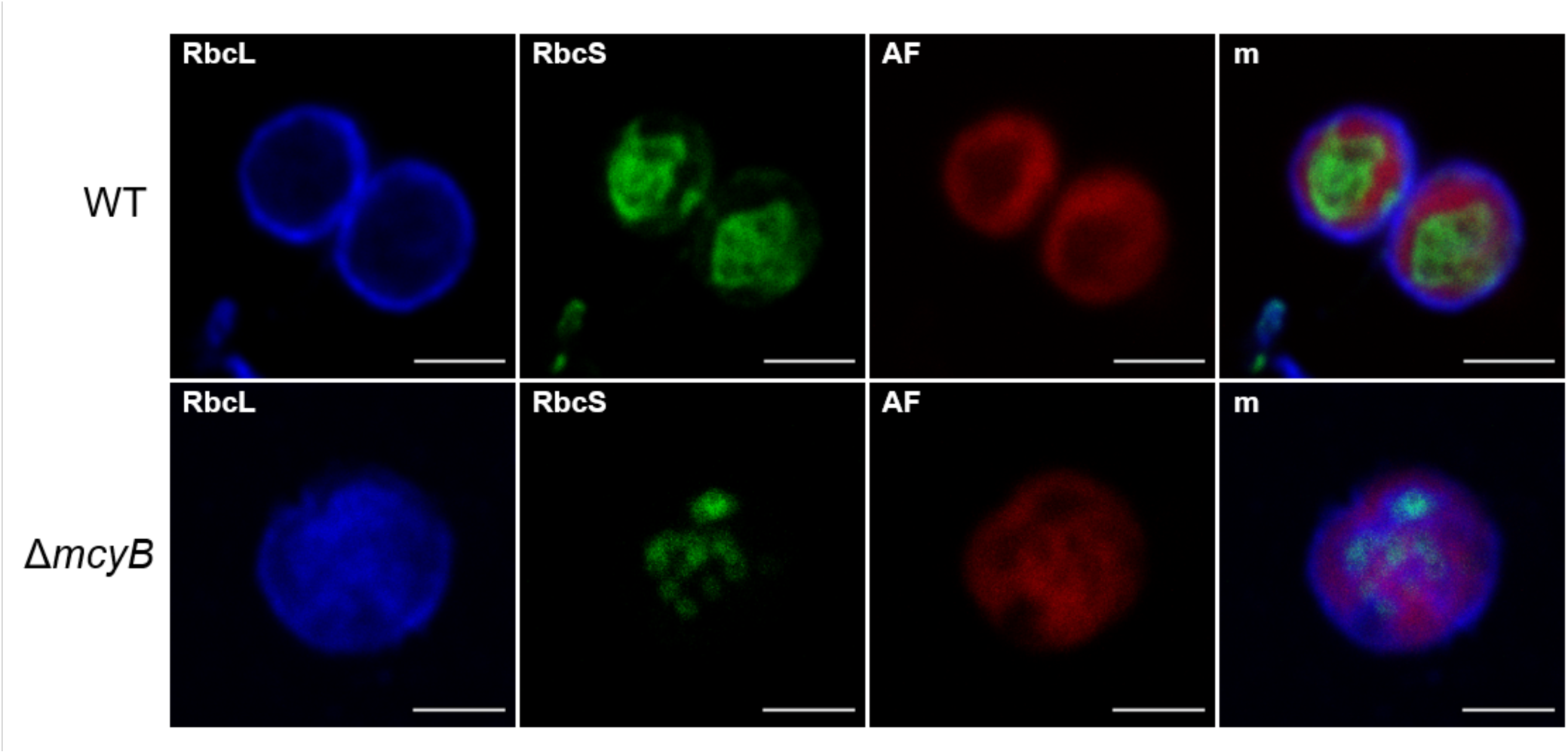
Co-Hybridization of high-light treated cells of **(A)** *M. aeruginosa* WT and **(B)** Δ*mcyB* with RbcL and RbcS antibodies (OD_750_: 0.6). RbcL is visible in the blue fluorescence channel and RbcS is visible in the green fluorescence channel. The fluorescence channel is indicated in the top left corner of each image. AF=phycobilisome auto fluorescence, m=merged image from the 3 fluorescence channels. The scale bar is 2 µm.

Cross-sections of low-light and high-light treated cells (OD_750_ 0.6) were also analyzed using transmission electron microscopy. Both WT and MC-deficient mutant cells showed a number of dark electron-dense granules (Fig. 6). While these granules were mostly scattered in the cytosolic and thylakoid areas in the MC-deficient mutant they were predominantly localized underneath the cytoplasmic membrane in the WT under high-light. Since their distribution pattern, their spot-like appearance and their light-dependent relocation closely resemble RbcL signals observed in IFM studies, we conclude that the granules may represent RubisCO-rich structures in *Microcystis*. At the same time, carboxysomes looked very pale, residing centrally in the cell without major redistribution. Only occasionally carboxysomes with a darker appearance, which has previously been reported in other cyanobacteria as being more characteristic for the electron-dense carboxysome structures, were observed (Fig. S3). Notably, electron-dense granules underneath the cytoplasmic membrane were recently also observed in an ultrastructural comparison of a toxic strain of *Microcystis* and a non-toxic strain of *Microcystis* (Jacinavicius et al. 2019). Taken together, subcellular fractionation studies, IFM and electron microscopic data all point to the same direction and suggest that i) the subcellular localization of RubisCO is dynamic in *Microcystis*, in particular after high-light treatment at high cell density; ii) a substantial fraction of RbcL is primarily located outside of carboxysomes; and iii) that a large portion of RbcL is associated with the cytoplasmic membrane.

**Fig. 6.**
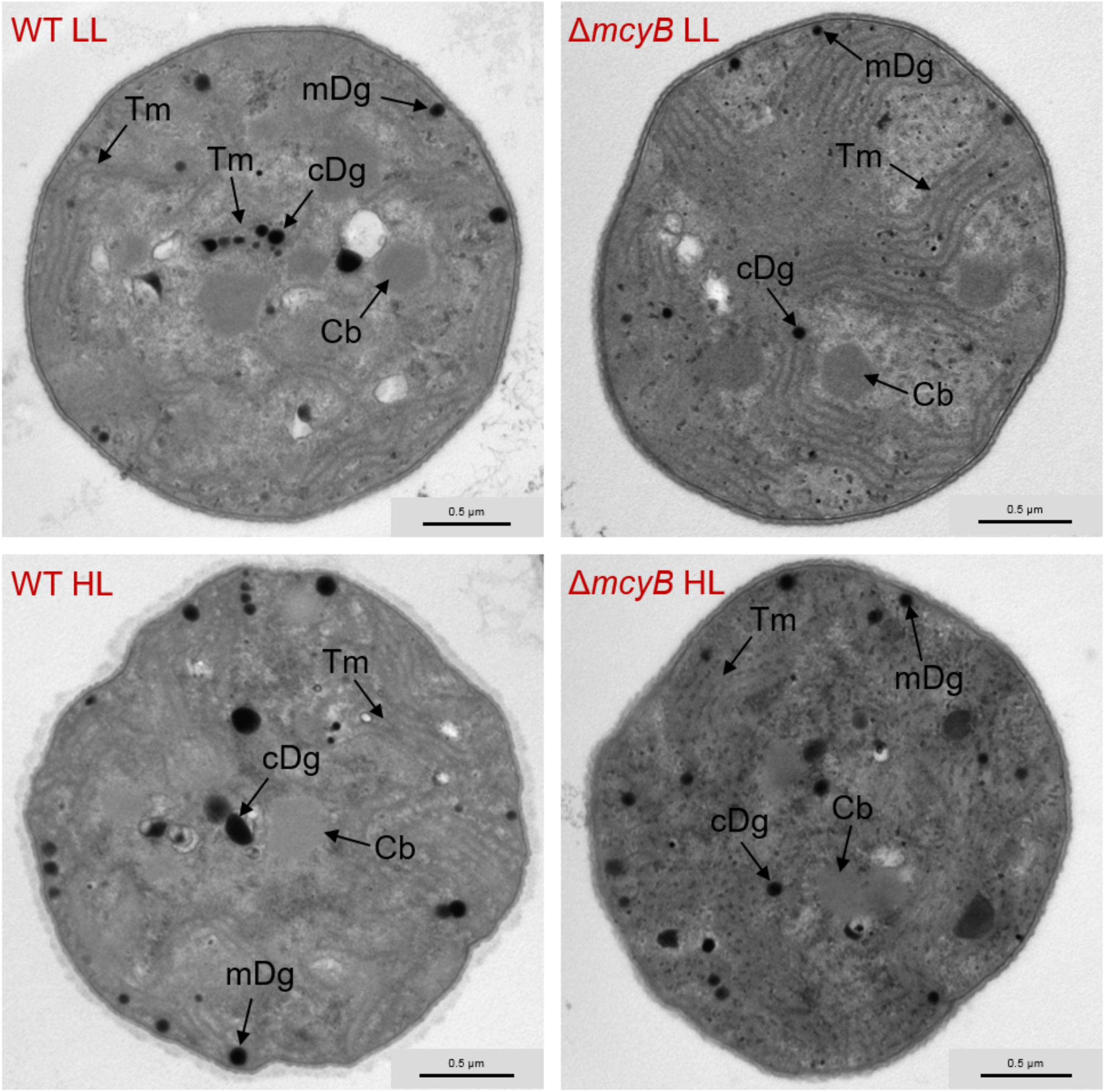
Transmission electron micrographs (TEM) of *M. aeruginosa* wild type (WT) and Δ*mcyB* mutant cells (MT) under low-light conditions and after 3 h of high-light treatment. The upper images display low-light (LL) adapted cells of WT and Δ*mcyB* and the lower images display cells after 3h of high-light (HL) treatment. Light conditions were as for Fig. 1. Electron dense granules are highlighted with black arrows. After 3h of high-light treatment more granules are located near the cytosolic membrane, especially in the WT. Carboxysomes appear very pale in both WT and Δ*mcyB* mutant. See Figure S3 for further examples. Tm: thylakoid membrane, Cb: carboxysomes, cDg: cytosolic dense granules, mDg: cytosolic membrane-associated dense granules.

### Subcellular localization of protein-bound microcystin

In order to assess possible spatial effects on RubisCO-MC interactions, we analyzed the subcellular localization of MC biosynthesis proteins and MC itself in WT cells under high-light conditions. The non-ribosomal peptide synthetase McyB and the aspartate racemase McyF were detected by IFM using specific antibodies against these proteins. To our surprise, both proteins were primarily located at the thylakoid membrane facing the small cytosolic space of *Microcystis* (Fig. 7A-D). We conclude that the MC biosynthesis complex is anchored to the thylakoid membrane. To visualize the protein-bound MC portion by IFM we utilized a commercially available antibody against MC. Microcystin was predominantly detected in small distinct spots in the cytosolic area and at the thylakoids but was also visible at underneath the cytoplasmic membrane (Fig. 7E-H). Control experiments without primary antibody did not yield any specific signal (Fig. S2). As the soluble MC pool has been largely lost during the fixation process the signals were primarily attributed to the protein-bound portion of MC. Co-localization studies of MC with RbcL and RbcS respectively, revealed a perfect overlap of distinct MC spots in the cytosol with RbcS-containing spots (Fig. 7I-L). RbcS and MC were also frequently co-located at the inner thylakoid membrane (Fig. 7M-P). In contrast, RbcL only co-located with MC at the cytoplasmic membrane (Fig. 7E-H). We hypothesize that *de novo* synthesized RbcS assembles near the MC biosynthesis machinery at the thylakoids and forms distinct RbcS species that separate from RbcL in *Microcystis*. Although MC was occasionally detected at the periphery of cells, we further conclude that RbcS rather than RbcL is a primary binding partner of MC.

**Fig. 7.**
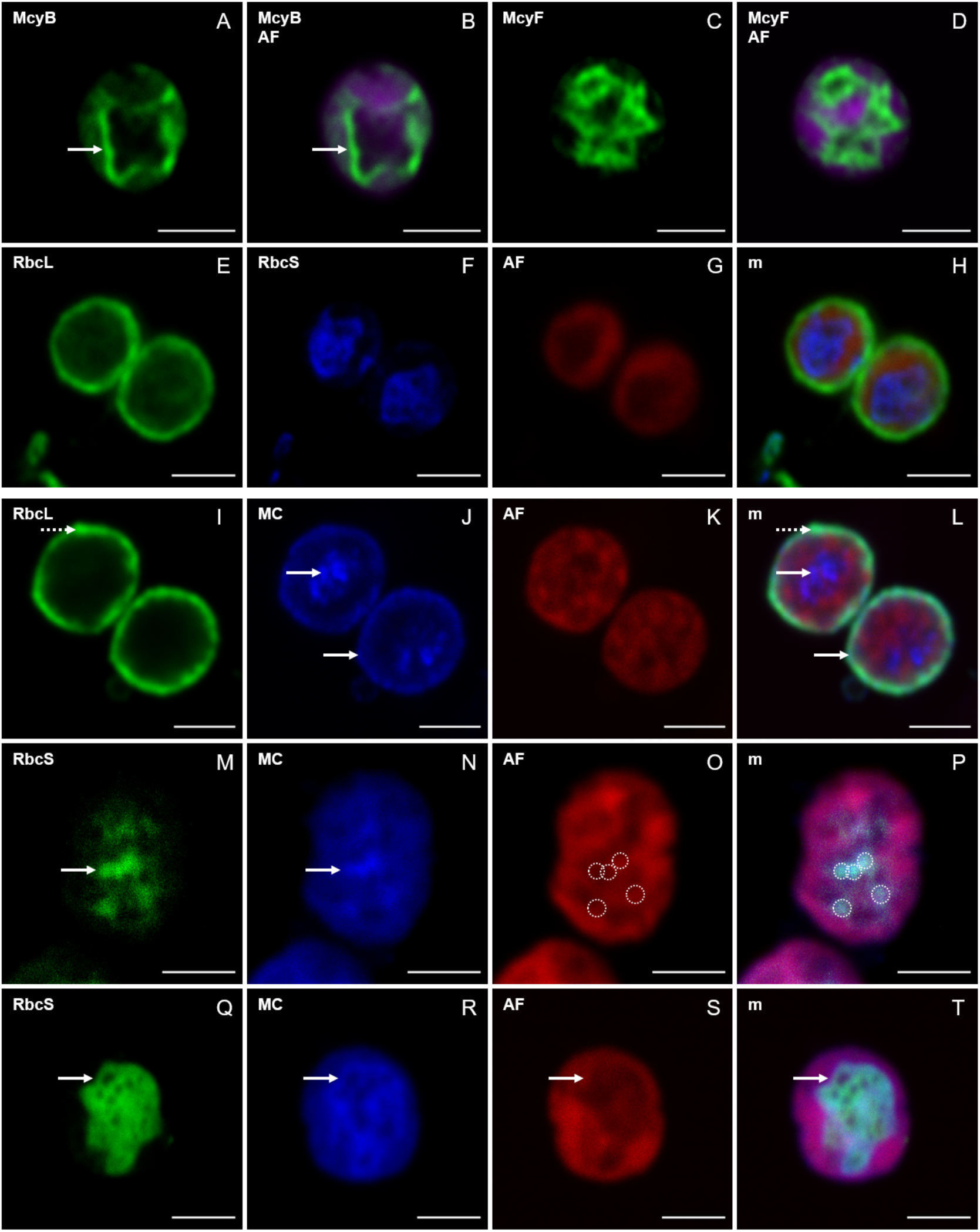
Location of microcystin biosynthesis proteins McyB and McyF and of protein-bound microcystin (MC) in high-light treated *M. aeruginosa* wild type visualized by IFM. **(A-B)** Location of McyB. The green fluorescence channel is displayed alone and merged with the phycobilisome autofluorescence (AF). The arrow indicates the strong signal of McyB located at the inner thylakoid membrane. **(C-D)** Location of McyF. The green fluorescence channel is displayed alone and merged with the chlorophyll autofluorescence (AF). McyF is located in spots at the inner thylakoid membrane. **(E-H)** Co-Localization of RbcL and MC. RbcL is shown in the green fluorescence channel and microcystin (MC) is shown in the blue fluorescence channel. The red channel shows the phycobilisome autofluorescence (AF). The dashed arrow indicates the localization of RbcL at the cytosolic membrane. MC mainly locates in the cytosol in spots (arrow). **(I-P)** Co-Localization of RbcS and MC. The fluorescence channel is indicated in the top left corner of each image. Both proteins co-localize strongly in several spots in the cytosol (I-P) and around the thylakoids (M-P). These spots are highlighted with dashed circles. AF: phycobilisome fluorescence, m: merged image from the 3 fluorescence channels. The scale bar in all images is 2 µm.

### RbcS and MC are part of a putative Calvin Benson cycle enzyme super complex

Next, we analyzed on native gels the existence of different RubisCO species in *M. aeruginosa* PCC7806, grown to high cell density (OD_750_: 0.6). High-molecular mass complexes comprising both RbcL and RbcS were observed within the soluble and the membrane fractions (Fig. 8A). A RubisCO complex of a similar size likely representing the canonical RbcL_8_S_8_ was also present in extracts of *Synechocystis* sp. PCC6803 and resembles the well-characterized RbcL_8_S_8_ complex reported for *Synechococcus elongatus* PCC6301 (Fig. 8A) (Liu et al., 2010). These data suggest that not only the cytosolic RubisCO but also the membrane-bound RubisCO of *M. aeruginosa* PCC7806 comprise a certain amount of RbcS, although the small subunit was not well detected at the cytoplasmic membrane *in situ*. However, RbcS was additionally detected in high-molecular mass complexes, which apparently do not contain the large subunit RbcL thereby further confirming the existence of separate RbcS species in *Microcystis.* Further separation of the membrane fraction from the MC-producing WT strain *M. aeruginosa* PCC7806 revealed that the RbcL-free RbcS containing complex remained tightly bound to the membrane even after repeated sonication steps, while membrane-bound RbcL_8_S_8_ could be easily detached from the membrane as described earlier (Fig. 8A, Fig. S1). The same blots from native PAGE were incubated with the anti-MC antibody. Here, we detected three major MC bands, one related to the putative RbcL_8_S_8_ complex and one related to the RbcL-free RbcS containing high molecular mass complex. A third band appeared that neither co-migrated with RbcL nor RbcS (Fig. 8A). Notably, MC-binding high-molecular mass complexes were not only present in the membrane fraction, where they comprised considerable amounts of RbcS, but also in the soluble fraction and the fraction loosely associated with membranes, where they did not comigrate with RbcL or RbcS (Fig. 8A).

**Fig. 8.**
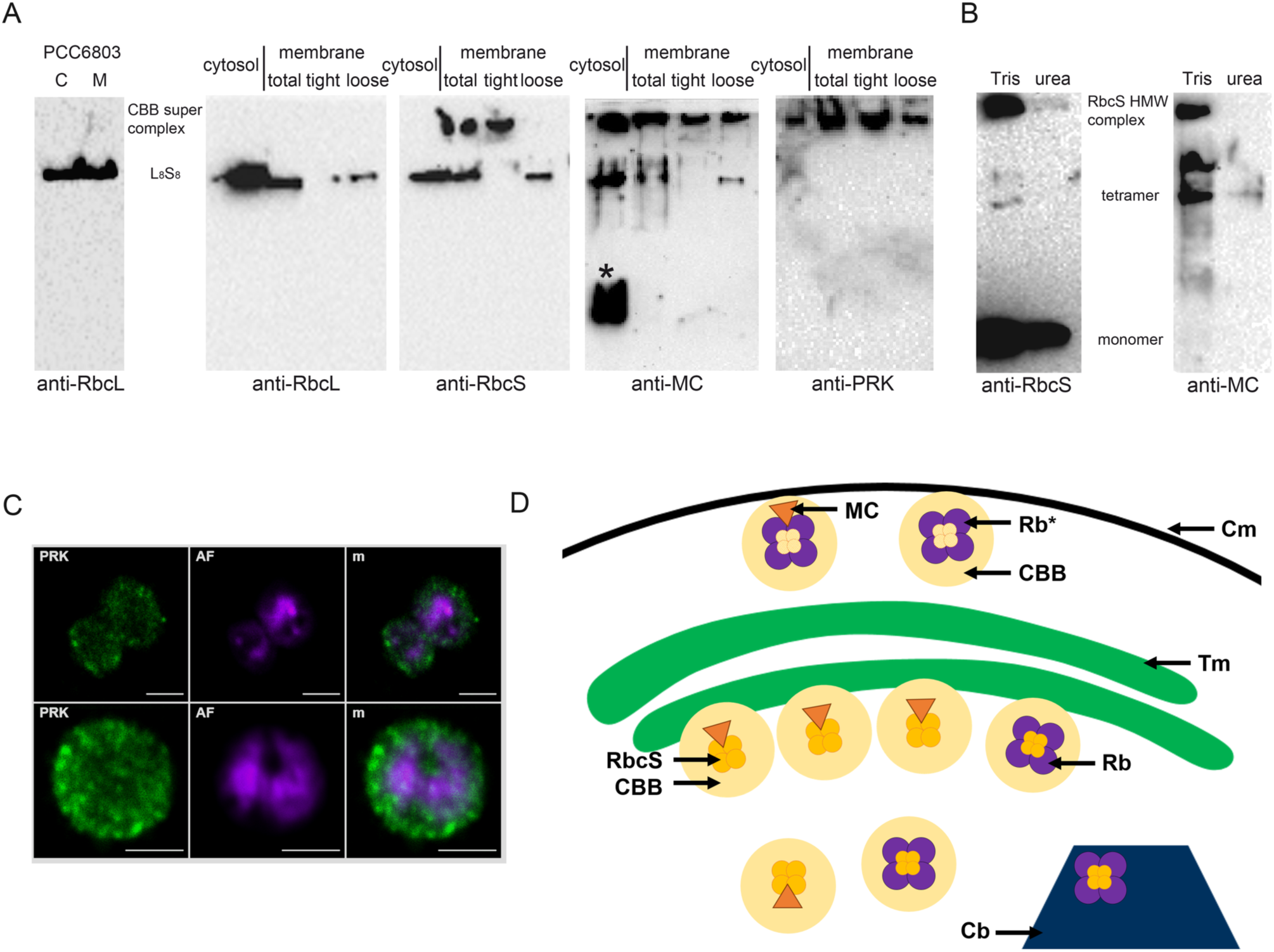
Native PAGE analysis of RubisCO complexes and RbcS-MC-binding analysis. **A)** Native Western blots indicating presence of RbcL_8_S_8_-like complex (RbcL_8_S_x_) in cytosolic and membrane protein extracts of *Synechocystis* sp. PCC6803 and *M. aeruginosa* PCC7806. The *Microcystis* extract additionally contains a RbcL-free RbcS-containing high-molecular mass complex. Total membrane fractions were further subfractionated into tightly bound proteins (tight) and loosely attached proteins (loose). Hybridization with anti-RbcL, anti-RbcS, anti-MC and anti-Prk is indicated below each blot. The MC binding signal not related to either RbcL or RbcS is indicated with an asterisk. **B)** Immunodetection of RbcS in RbcL-free RbcS fraction purified by anion exchange chromatography reveals presence of RbcS monomer along with SDS-stable RbcS oligomers and a RbcS high molecular weight complex (HMW). See Fig. S4 for further analysis of RbcS oligomers in thylakoid membrane preparations. Only the RbcS monomer is stable after treatment with 6M urea. Immunodetection of MC shows colocalization of MC signal with oligomeric forms of RbcS but not the RbcS monomer. MC signals disappeared after treatment with 6M urea. D) Schematic representation of soluble and membrane-bound RubisCO (Rb) and Calvin-Benson cycle super complexes (CBB). Rb* depicts RubisCO complexes at the cytoplasmic membrane that may contain lower amounts of RbcS according to IFM analysis. The orange triangle represents MC. The Calvin-Benson cycle super complex is shown as yellow cycle. Tm; thylakoid membrane, Cb; carboxysomes, Cm: cytoplasmic membrane.

Two independent proteomic studies have previously identified a striking number of Calvin-Benson cycle enzymes as predominant MC-binding partners (Wei et al., 2016; Zilliges et al., 2011). Hence, we considered the possibility that the high-molecular mass complexes comprising MC and RbcS represent Calvin-Benson cycle enzyme super complexes that are known from chloroplasts of eukaryotic algae and plants, which were also postulated for cyanobacteria (Agarwal et al., 2009; Suss, Arkona, Manteuffel, & Adler, 1993). To test this possibility, additional blots from native PAGE were incubated with an antibody directed against phosphoribulokinase (PRK). PRK was exclusively present detected in high-molecular mass complexes, and the signals exactly correspond to signals obtained with the anti-MC antibody. We thus postulate that PRK and MC may be part of a Calvin-Benson cycle enzyme super complex that putatively comprises all MC-bound Calvin-Benson cycle enzymes previously identified, which is present both in the soluble and the membrane fraction. This finding prompted us to test the subcellular localization of PRK using IFM (Fig. 8C). PRK was detected in spots in the cytosol and near the thylakoids but also showed a spot-like appearance underneath the cytoplasmic membrane. Purification of RbcL-free RbcS protein fractions by anion exchange chromatography from high-density cultures of *M. aeruginosa* PCC7806 WT revealed a number of SDS-stable oligomers and a high-molecular mass complex along with monomers of RbcS (Fig. 8B and Fig. S4). Furthermore, RcbL-free RbcS fractions of approximately 55 kDa likely representing stable tetramers of RbcS were also detected in thylakoid membrane preparations of *M. aeruginosa* PCC7806 (Fig. S4). The oligomeric/higher molecular mass forms of RbcS disappeared under denaturing conditions, where only the RbcS monomers remained detectable (Fig. 8C and S5D). This finding also excludes a possible cross-reaction with the 35 kDa or 58 kDa form of the carboxysome protein CcmM that contains a RbcS-like domain. The same fractions were tested with the MC antibody that yielded strong signals with different oligomeric/high molecular mass forms of RbcS but not the monomeric form. The microcystinylation signals largely disappeared under denaturing conditions (Fig. 8C). These data suggest that i) MC specifically binds to RbcS oligomers and RbcS containing high-molecular mass complexes; and ii) that the interaction between RbcS and MC is noncovalent

## Discussion

Much of the current view on the primary metabolism of cyanobacteria and on RubisCO is derived from studies on a few unicellular model strains like *Synechocystis* sp. PCC6803 and *S. elongatus* PCC7942 (Cameron, Wilson, Bernstein, & Kerfeld, 2013; Marcus, Altman-Gueta, Finkler, & Gurevitz, 2003). Research on cyanobacteria featuring more complex life styles is largely hampered by the lack of molecular techniques and the difficulty to grow and keep the cyanobacteria in an axenic state. In spite of its high ecological relevance *M. aeruginosa* PCC7806 is a representative of this unexplored majority. The data presented herein unequivocally suggest that a substantial part of RubisCO is found outside the carboxysomes and is mainly located underneath the cellular membrane in high-light treated cells of the *Microcysti*s WT. Furthermore, the widespread cyanotoxin MC contributes to the versatility of RubisCO in *Microcystis* by interfering with the subcellular localization, its membrane association and assembly dynamics *in vivo*.

While single-celled cultures of *M. aeruginosa* PCC7806 grown under low light and ambient air show an apparent cytoplasmatic and carboxysomal localization of RubisCO at low cell density, a pronounced dynamic in the localization of RubisCO was observed at higher cell densities. One may therefore speculate that the phenotypic heterogeneity observed for RubisCO reflects the multicellular lifestyle of this colonial cyanobacterium, while the low density single-celled state may more closely resemble planktonic unicellular cyanobacteria such as *Synechococcus* and *Synechocystis* sp. MC may be one of the specialized molecules that differentiate low-density cultures from high-density cultures as the toxin specifically associates with protein complexes and is increasingly secreted as a signal at higher cell density (Wei et al., 2016). Phenotypic heterogeneity of multicellular cyanobacteria with regard to carbon fixation activities was already reported in several field studies utilizing the high-resolutions NanoSIMS technology for the measurement of ^13^C fixation rates. Foster et al. have compared carbon fixation activities in single-celled and colonial types of *Crocosphera watsonii* and observed major differences in the amount of assimilated carbon among cells within colonies but not with single cells (Foster, Sztejrenszus, & Kuypers, 2013). Furthermore, a recent NanoSIMS analysis of the filamentous bloom-forming cyanobacterium *Nodularia spumigena* that is producing the MC-like peptide nodularin indicated a heterogeneity in carbon fixation activities at the ultrastructural level within single filaments (Schoffelen et al., 2018). Both studies reported that carbon fixation activities are often concentrated in small spots resembling the membrane-associated granules observed in our electron microscopic studies (Foster et al., 2013; Schoffelen et al., 2018). A study on *Anabaena oscillaroides*, again, has even demonstrated a predominant CO_2_ fixation activity in spots underneath the cytoplasmic membrane suggesting that other cyanobacteria than *Microcystis* may also concentrate RubisCO carbon-fixation activities at the periphery of cells (Popa et al., 2007). Collectively, these studies suggest that the CCM may not solely rely on RubisCO in carboxysomes in these multicellular collectives. The assembly process of both the hexadecameric RubisCO complex and of carboxysomes, respectively, is rather complex (Cameron et al., 2013; Liu et al., 2010) and lacks the dynamics experienced in cyanobacterial blooms (CO_2_, O_2_, light). Cytoplasmic membrane localization of RubisCO as observed during this study may thus be of particular relevance for short term high-light episodes that lead to a fast *de novo* synthesis of RubisCO.

Membrane localization of amphitropic proteins such as RubisCO can affect functions of proteins in different ways: i) through a closer localization with the substrate, activator or downstream target, or ii) through activation of the protein by a conformational switch (Johnson & Cornell, 1999). We can only speculate whether cytoplasmic membrane localization of RubisCO in *Microcystis* may indeed facilitate a closer interaction with the substrate CO_2_. Sandrini *et al.* (13) have predicted the existence of two homologs of the periplasmatic carbonic anhydrases (CA) EcaA and EcaB in all strains of *Microcystis* that could potentially enrich CO_2_ in the periplasm, although experimental evidence is currently missing. A high abundance and activity of extracellular EcaA and EcaB homologues was recently demonstrated for *Cyanothece* ATCC 51142 (Kupriyanova et al., 2019). In the field, CO_2_ may also be provided due to the respiratory activity by heterotrophic bacteria that are intimately associated with *Microcystis* and promote its growth. The possible CO_2_ enrichment in the periplasm may also explain that strain *M. aeruginosa* PCC7806 can cope without the high affinity bicarbonate uptake transporter SbtA that forms a pivotal part of the canonical CCM in other cyanobacteria. EcaB may additionally reside in the thylakoids as demonstrated in *Synechocystis* sp. PCC6803 (Sun et al., 2019). The closer proximity to carbonic anhydrase activities outside the carboxysome may thus partly explain the higher accumulation of CO_2_ fixation products observed in the *M. aeruginosa* PCC7806 WT *in vivo* (Fig. 1).

Our study further provides evidence for a close connection of RubisCO with a membrane-bound and a soluble Calvin-Benson cycle enzyme super complex further converting the RubisCO carbon-fixation products. As PRK was also detected underneath the cytoplasmic membrane (Fig. S4), the Calvin-Benson cycle is likely also intimately associated with RubisCO residing at the cytoplasmic membrane. The advantage of juxtaposing sequentially acting enzymes in proximity of membranes that limit unwanted diffusion of intermediates in a hydrophobic environment has been discussed earlier (Agarwal et al., 2009; Suss et al., 1993). The Calvin-Benson cycle can further capitalize on the proximity of ATP synthase that is also residing in the thylakoids and represents another proven MC-binding partner (Wei et al., 2016). A joint localization of ATP synthase and Calvin-Benson cycle enzymes has been observed in other cyanobacteria before (Agarwal et al., 2009).

In *Microcystis*, however, the neighbourhood is expanded by the MC biosynthesis complex and MC itself. We hypothesize that the McyH ABC transporter featuring a membrane domain may serve as a membrane scaffold (Pearson, Hisbergues, Borner, Dittmann, & Neilan, 2004). MC production has a high demand of ATP and can strongly benefit from the proximity to ATP synthesis. MC, in turn, can stabilize Calvin-Benson cycle enzymes and promote binding of RbcS to the membrane and granule formation. Thylakoid-bound RbcS may serve as pool that can reversibly join with RbcL as seen in the dark phase of our diurnal experiment. MC production is clearly enhanced under high-light, however, the excess MC produced under these conditions immediately binds to proteins and does not appear in the free MC pool (Meissner et al., 2013). We are thus speculating that MC production, RubisCO delocalization, RbcS oligomerization and Calvin cycle super complex formation are closely connected and tether the complex to either the thylakoid or cytoplasmic membrane. Because blocking of cysteines prevents microcystinylation *in vitro*, we have previously postulated that MC-binding to proteins in *Microcystis* is facilitated by covalent binding to cysteines. Our new data rather suggest that blocking of cysteines prevents formation of oligomeric forms of RbcS and thus indirectly prevents interactions with MC (Zilliges et al., 2011).

Considering the major differences in the accumulation of RubisCO carboxylation products in WT and Δ*mcyB* mutant, it seems surprising that other *Microcystis* strains have selectively lost the capability to produce these toxins. Non-MC-producing *Microcystis* strains, however, produce other peptides instead including amphitropic molecules like microginins and aeruginoguanidines (Makower et al., 2015; Pancrace et al., 2018). We speculate that the different peptides further contribute to the previously described genotypic and phenotypic heterogeneity in the CCM of *Microcystis*. Heterogeneity in the response to fluctuating C_i_ and light conditions may be beneficial at the community level. A flexible CCM mechanism may also contribute to the robustness of growth when multicellular *Microcystis* colonies experience C_i_ limitation especially under high-light conditions. In agreement with this hypothesis, Paerl *et al*. have demonstrated a partitioning of carbon fixation when freshly isolated *Microcystis* colonies were exposed to C_i_-limiting conditions (Paerl, 1983).

Our data strongly suggest that RubisCO in *Microcystis* is more versatile than previously expected. Yet, the low-light to high-light shift experiments applied in our study provide only a little snapshot into the RubisCO localization and activities. One may speculate that secreted levels of 2-PG as detected during this study have a much higher relevance under the extreme high-light and oxygen conditions experienced in *Microcystis* blooms at the surface of lakes. *Microcystis* colonies could thereby fuel their heterotrophic microbiome in an advanced mutualistic relationship and capitalize on the bifunctionality of RubisCO. Positioning of RubisCO at the membrane is likely just one aspect of a larger network of metabolic facilitation between bloom-forming cyanobacteria and their heterotrophic microbiomes. We also need to explore the structural basis for the interaction of MC and RbcS oligomers. The present study sets the ground to further understand structure-activity relationships and highlights the importance of considering the membrane localization of RubisCO for the interpretation of *in vivo* metabolomic data.

## Material and Methods

### Cultivation conditions

*Microcystis aeruginosa* PCC7806 was cultivated in liquid BG-11 medium (Rippka, 1988). Chloramphenicol in a final concentration of 3 µg/ml was added to BG-11 for cultivation of the Δ*mcyB* mutant (Dittmann, Neilan, Erhard, von Dohren, & Borner, 1997). The strains were grown at 23°C under continuous illumination at 10 µmol photons m^-2^ s^-1^ without agitation or external aeration to obtain low light adapted cultures. Growth was monitored by measuring the optical density at 750 nm. For high light experiments the WT and the Δ*mcyB* mutant cultures were diluted with BG-11 to an OD_750_ of 0.2 and incubated at low light conditions until an OD_750_ of 0.45 was reached. Subsequently, the cultures were divided into 4×80 ml and were then transferred into a Multi-Cultivator device (MC 1000, Photo Systems Instruments). The cultures were illuminated at a light intensity of 250 µmol photons m^-2^ s^-1^ (high light) for 4 h under continuous aeriation with ambient air. 50 ml of sample was taken at the start of the experiment and every hour of the 4 h high light illumination for protein extraction and microscopy. The sample used for protein extraction was centrifuged at 21,000 x *g* for 7 min at 4°C and the pellet was stored at −20°C for later extraction. The growth of the cultures was monitored by measuring the optical density at 750 nm at every sampling point. The experiment was repeated 3 times with similar results. For the diurnal experiment, cultures of *M. aeruginosa* WT and the Δ*mcyB* mutant were cultivated under a 16h light (60 µmol photons m^-2^ s^-1^), 8h dark cycle. Samples were taken after the cultures reached an OD_750_ of 0.3 (low cell density experiments) or 0.6 (high cell density experiments) during the day and the night, respectively.

### Subcellular Fractionation

All the following steps were performed on ice or pre-cooled centrifuges. The cell pellet was re-suspended in 500 µl Native Extraction Buffer (50 mM HEPES; 5mM MgCl_2_ x 6 H_2_O; 25 mM CaCl_2_ x 2 H_2_O; 10 % glycerol; pH set to 7.0) in a 1.5 ml reaction tube. The sample was sonicated (Sonopuls mini20, Bandelin) for 90 secs (50 % amplitude, 3 secs on/off) and PMSF was added at a final concentration of 1 mM. Subsequently, a slow centrifugation (2,000 x *g* for 2 min) was performed to get rid of unbroken cells, followed by a long centrifugation of the supernatant (21,000 x *g* for 15 min). The resulting supernatant was transferred into a new reaction tube (cytosolic fraction) and the pellet was re-suspended again in 500 µl Native Extraction Buffer. Half of the volume (250 µl) was saved as the total membrane fraction (membrane-associated proteins) and stored at −20°C. The remaining sample was sonicated and centrifuged with the same parameters as the first time (without the slow centrifugation step), the supernatant is the loose fraction (loosely attached proteins to membranes). The pellet was re-suspended in 250 µl Native Extraction Buffer and is the tight fraction (tightly bound proteins to membranes). All protein samples were stored at - 20 °C until use. For the extraction of the total membrane protein fraction, the pellet of the first sonication step was re-suspended in Urea Buffer (8 M Urea; 100 mM NaH_2_PO_4_; 100 mM Tris-HCl; pH set to 8) instead of Native Extraction Buffer. After sonication the sample was stored at −20°C without any further centrifugation (membrane fraction).

Thylakoid membrane extraction, running of the blue-native PAGE and preparation of the gel for 2^nd^ dimension SDS-PAGE were performed as described previously (Gandini, Schmidt, Husted, Schneider, & Leister, 2017).

### Heterologous expression of proteins and antibody generation

Following primers were used for PCR-amplification of *rbcS* (IPF_2530) and *ccmk2* (IPF_5495) from *Microcystis aeruginosa* PCC 7806 genomic DNA: 5′-ttt<u>catATG</u>AAAACTTTACCTAAAGAGAAGCGTTA-3′ and 5′-ttt<u>ggatcc</u>TTAGTAGCGGCCGGCATTG-3′ for *rbcS* and 5′-ttt<u>catATG</u>CCAATTGCAGTAGGAATGA-3′ and 5′-ttt<u>ggatcc</u>TTAATAGCTGCGGAATTGCT-3′ for *ccmk2*. Using *NdeI* and *BamHI*, gel-purified PCR products were ligated into the pET15b expression vector (Novagen). Expression in *E. coli* BL21 (DE3) cells (Novagen) was induced with 0.5 mM Isopropyl β-D-1-thiogalactopyranoside at OD_600_ of ∼1.0, cells were grown for 2 h at 37°C with shaking at 220 rpm. Cell pellets were resuspended in denaturing buffer (100 mM Na_2_HPO_4_; 10 mM Tris-HCl; 8 M urea; 10 mM imidazole; pH 8.00) and lysed by sonication. His-tagged proteins were purified over Ni-NTA-agarose (Qiagen) with 3 wash steps at 30 mM imidazole and elution with 250 mM imidazole in denaturing buffer. Protein purity was determined by SDS-PAGE, 1 mg of denatured protein was used to raise polyclonal antibodies in rabbit serum (Pineda antibody service, Berlin, Germany).

### SDS-PAGE and immunoblotting

Proteins were separated by Bis-Tris SDS-PAGE (recipe from BiteSize Bio based on NuPAGE from Invitrogen) on a polyacrylamide gel (8-15 %, depending on the targeted protein). Total protein extracts were loaded on each lane and the gel was run at constant voltage of 190 V for 35 min. To detect the targeted protein, they were blotted on a nitrocellulose membrane (Protein Premium 0.45 µm MC; Amersham) as described previously (Towbin et *al*., 1979). The transfer buffer contained 20 % methanol for a more efficient blotting. After blotting the membrane was blocked with 1 % polyvinylpyrrolidone (PVP) K-30 in TBS-T (6.06 g/l Tris; 8.77 g/l NaCl; pH set to 7.4; add 0.1 % Tween-20) and was washed subsequently at 4°C. The primary antibodies were incubated in TBS-T with the following dilutions overnight at 4°C: RbcL 1:10000; RbcS 1:5000; CcmK 1:5000; FtsZ 1:5000; MC 1:5000; PRK 1:5000. The membrane was washed with TBS-T to remove unbound primary antibody and the secondary antibody (α-Mouse-IgG HRP-conjugate for MC and α-Rabbit-IgG HRP-conjugate for the remaining antibodies) was applied to the membrane at a dilution of 1:10000 in TBS-T and incubated at 4°C for at least 1 h. Afterwards, the membrane was washed 4 times and developed (SERVALight Polaris CL HRP WB Substrate Kit, Serva). Images were taken with the ChemiDoc XRS+ Imaging System (Bio-Rad). It is of note that we observed SDS-stable oligomers of RbcS and CcmK in both WT and mutant protein fractions. In order to fully quantify the two proteins, the corresponding protein gels were thus treated with 4M urea (Fig. 2B). We did not observe cross-reactivity of the antibody with the 58 kDa or 35 kDa form of the carboxysome shell protein CcmM that contains a RbcS domain in our Western Blot experiments.

### Immunofluorescence microscopy (IFM)

4 ml of culture were separated into two 2 ml reaction tubes and centrifuged 1 min at 10000 x *g*, 4°C (condition for every centrifugation step). To wash the cells, the pellet from one tube was re-suspended with 1 ml of phosphate-buffered saline (PBS: 8.18 g/l NaCl; 0.2 g/l KCl; 1.42 g/l Na_2_HPO_4_: 0.25 g/l KH_2_HPO_4_; pH set to 8.3), transferred to the other tube to resuspend the pellet as well and was centrifuged again. For fixation the pellet was resuspended with 1 ml of 4% formaldehyde in PBS. The *M. aeruginosa* WT samples were incubated for 30 min, the Δ*mcyB* mutant for 15 min at room temperature. After two washing steps with PBS the pellet was re-suspended with 100 µl of water and 20 µl were spread on a microscope slide each. The slides were air-dried and stored at −20°C for later use.

To start the hybridization with antibodies the sample slides were equilibrated in PBS for 5 min at room temperature. Afterwards, the slides were incubated with 2 mg/ml lysozyme in PBS-TX (PBS with 0.3 % Triton X-100) for 30 min at room temperature and washed twice with PBS-TX for 3 min. The samples were blocked with 1% PVP K-30 in PBS-T (PBS with 0.3 % Tween-20) for at least 1 h at 4°C and washed twice with PBS-T. The primary antibody dilutions were made in PBS-T as well: RbcL 1:300 Rubisco large subunit form I, chicken; Agrisera); RbcS 1:200 (rabbit); CcmK 1:200 (rabbit); MC 1:250 (Microcystin-LR, mouse; Enzo Life Sciences); McyB 1:100 (rabbit); McyF 1:100 (rabbit); PRK 1:200 (rabbit). After an incubation for at least 1 h at 4°C the slides were washed twice to remove unbound primary antibody and the secondary antibody was applied to the slides. Alexa Fluor 488 goat anti-rabbit (1:200), Alexa Fluor 488 goat anti-mouse (1:100), Alexa Fluor 546 goat anti-chicken (1:200) and Alexa Fluor 568 goat anti-mouse (1:100) were used as secondary antibodies depending on the selected primary antibodies. Subsequently, the slides were washed, air-dried and a drop of glycerol containing 4% propyl gallate was applied to the slide and covered with a coverslip. The slides were stored at −20°C until use. Immunofluorescence images were taken with a Zeiss LSM 780 laser scanning confocal microscope using a Plan-Apochromat 63x/1.40 oil immersion objective. Alexa Fluor 488 was excited at 488 nm (detection spectrum 493 – 556 nm), Alexa Fluor 546 and 568 at 561 nm (570 – 632 nm), and autofluorescence at 633 nm (647 – 721 nm). The excitation was performed simultaneously.

### Electron microscopy

*M. aeruginosa* PCC7806 WT and Δ*mcyB* cells were diluted with fresh BG-11 medium and grown under low light conditions until an OD_750_ of 0.4 was reached. Then 50 ml of the cultures were irradiated with high-light (250 µmol photons m^-2^s^-1^) for 3 h. 2 ml samples were taken before (t_0_) and after 3 h high-light exposure (t_3_). The samples were centrifuged for 2 min at 13,000 x *g*, the supernatant was removed, and the fixative (2.5% glutaraldehyde, 2.0% formaldehyde in 0.1 M Na-cacodylate buffer, pH 7.4) was added directly on the pellet without re-suspension. Samples were fixed for 1 to 3 h at room temperature, then over night at 4°C. Samples were washed 3 times for 10 min in 0.1 M Na-cacodylate buffer and post-fixed for 90 min at room temperature in Na-cacodylate-buffered 2% OsO_4_. After washing twice for 10 min in H_2_O, samples were overlaid by a thin layer of 1% low-melting agarose, dehydrated in a graded EtOH series and acetone, and embedded in low viscosity resin (Agar Scientific Ltd., Stansted, Essex, UK). Ultrathin sections stained with uranyl acetate and lead citrate were examined in a JEM 1011 (JEOL Ltd., Tokyo, Japan).

### LC/MS sample preparation and measurement

For analysis of metabolites three separate cultures of *M. aeruginosa* PCC7806 WT and Δ*mcyB* were grown under low light conditions until an OD_750_ of 0.38 was reached. 80 ml of culture were radiated with high-light (250 µmol photons m^-2^s^-1^) for 4 h. At the start of irradiation and every following hour a sample of 7.5 ml was taken and centrifuged for 7 min, 21,000 x *g* at 4°C. The supernatant was filter sterilized (0.45 µm), frozen with liquid nitrogen and stored at −20°C for later use (extracellular metabolites fraction). The pellet was frozen with liquid nitrogen as well and stored at −20°C. For extraction of the metabolites the pellet was re-suspended with 4 ml H_2_O and sonicated for 2 min (60%, 3 secs on/off). After centrifugation for 10 min, 21,000 x *g* at 4°C the supernatant (intracellular metabolites) and the previously stored extracellular metabolite fractions were dried in a vacuum concentrator (RVC 2-25 CDplus, Christ). The dried extracts were dissolved in 200 µl water and filtrated through 0.2 µm filters (Omnifix®-F, Braun, Germany). The cleared supernatants were analyzed using the high-performance liquid chromatograph mass spectrometer LCMS-8050 system (Shimadzu, Japan) and the incorporated LC-MS/MS method package for primary metabolites (version 2, Shimadzu, Japan). In brief, 4 µl of each extract was separated on a pentafluorophenylpropyl (PFPP) column (Supelco Discovery HS FS, 3 µm, 150 × 2.1 mm) with a mobile phase containing 0.1% formic acid. The compounds were eluted at 0.25 ml min^-1^ using the following gradient: 1 min 0.1% formic acid, 95% *A. dest*., 5% acetonitrile, within 15 min linear gradient to 0.1% formic acid, 5% A. dest., 95% acetonitrile, 10 min 0.1% formic acid, 5% *A. dest*., 95% acetonitrile. Aliquots were continuously injected in the MS/MS part and ionized via electrospray ionization (ESI). The compounds were identified and quantified using the multiple reaction monitoring (MRM) values given in the LC-MS/MS method package and the LabSolutions software package (Shimadzu, Japan). Authentic standard substances (Sigma-Aldrich, Germany) at varying concentrations were included in all batches and used for calibration.

### Rubisco purification from *Microcystis aeruginosa* PCC 7806

Intact, functional Rubisco was purified from *Microcystis aeruginosa* PCC 7806 by fractionated ammonium sulfate precipitation coupled with anion exchange FPLC (adapted from (Salvucci, Portis, & Ogren, 1986)). After centrifugation of 600 mL of liquid culture, cell pellets were washed with ice-cold water and shock frozen in liquid nitrogen. Pellets were thawed on ice and resuspended in 40 mL of ice-cold Rubisco extraction buffer (10 mM Bicine, 1 mM EDTA, 1 mM DTT, pH 8.1). Cells were lysed with a cell disruptor (T-series, Constant systems Ltd) at 35 kPSI. DNA was sheared by sonication for 5 min, cell debris was pelleted at 20,000 x *g*, 4°C for 1h. Hydrophobic proteins were precipitated for 30 min at 4°C with light shaking after a 20% saturated solution of ammonium sulfate was generated by the slow addition of a fully saturated ammonium sulfate solution. After centrifugation at 20,000 g, 4°C, 10 min, the supernatant was adjusted to 50% saturated ammonium sulfate and kept shaking at 4°C at least 1 h or overnight. The remaining proteins (including Rubisco) were pelleted (20,000 x *g*, 4°C, 10 min) and any residual supernatant was removed completely. Pellets were resuspended in 15 mL Rubisco extraction buffer + 0.5% w/v Triton X-100 and incubated at 4°C, 1 h with light shaking to solubilize any remaining lipid membrane patches. To remove residual ammonium sulfate and to precipitate Rubisco and other soluble proteins, PEG6000 was added to a final concentration of 20% w/v and the solution was incubated as before. Proteins were pelleted, the supernatant was removed completely as before and protein pellets were resuspended in 10 mL of buffer A (100 mM K_2_HPO_4_, 1 mM EDTA, 1mM DTT, pH 7.6).

The resulting suspension was cleared by centrifugation and filtered through a 0.45 µm syringe filter. The sample was loaded on a MonoQ 4.6/100 PE anion exchange column run on an ÄKTApurifier FPLC system (both GE Healthcare) with following parameters: flow rate: 0.2 mL/min; equilibrated with 5 CV; flowthrough fractionation: 2 mL; empty 5 mL sample loop with 10 mL; wash out unbound sample: 2 CV; eluate fraction size: 2 mL; linear gradient; target conc buffer B: 50%; length of gradient: 50 CV; gradient delay: 2 mL; clean after elution: 15 CV. Buffer B was 1 M KCl, 100 mM K_2_HPO_4_, 1 mM EDTA, 1mM DTT, pH 7.6. Rubisco typically appeared as a distinct peak after ca. 55 mL elution volume at around 35% B. Purity was assessed with SDS PAGE and anti-RbcL (AS03 037A, Agrisera) and anti-RbcS immunoblots.

### Rubisco activity measurements

Purified Rubisco was assayed for carboxylase activity essentially as in (Parry et al., 1997). Rubisco was activated for 5 min at 25°C in 20 mM Bicine, 50 mM MgCl_2_, 50 mM NaHCO_3_ (pH 8.0) and then diluted to final concentrations between 0.02 mg/mL and 0.05 mg/mL into assay buffer containing 100 mM Bicine, 20 mM MgCl_2_, 30 mM NaHCO_3_, 1 mM NaH^14^CO_3_ (pH 8.1). The reaction was initiated by the addition of Ribulose-1,5-bisphosphate (RuBP) to a final concentration of 0.35 mM in a total assay volume of 300 µL. The reaction was stopped at varying intervals within a time frame of 10 min by transferring a 50 µL assay aliquot to 200 µL of 10 M formic acid. This sample was then completely evaporated at 80°C and the remaining pellet was resuspended in 500 µL water. To this, 5 mL of a liquid scintialltion cocktail (Ultima Gold, PerkinElmer) were added and the samples analyzed in a scintillation counter (Tri-Carb 2810TS, PerkinElmer). Specific carboxylase activities were calculated as nmol of fixed ^14^C per minute and mg Rubisco.

## Author contributions

E.D. and M.H. designed research, T.B., A.G., S.M., S.T., M.H., O.B. performed research. E.D. wrote the paper with contributions from all authors.

## Acknowledgements

The work was supported by grants from the Deutsche Forschungsgemeinschaft (DFG) to ED (Di910/10-1) and MH (Ha2002/20-1) and the DFG-funded Collaborative Research Centre ChemBioSys (SFB 1127) to ED. The LC-MS equipment at University of Rostock was financed through the HBFG program (GZ: INST 264/125-1 FUGG).

## Competing financial interests

The authors declare no competing financial interests.

