## Supplemental Figures 1-4 for "Non-canonical localization of RubisCO under high light conditions in the toxic cyanobacterium *Microcystis aeruginosa* PCC7806"

**Figure S1** RbcL is loosely attached to membranes in *M. aeruginosa* PCC7806 and the ∆*mcyB* mutant.

**Figure S2** Control immunofluorescence hybridizations with secondary antibodies used for detection of RbcL and CcmK.

**Figure S3** Transmission electron micrograph of *M. aeruginosa* PCC7806 indicating presence of carboxysome structures with differences in electron densities

**Figure S4** Blue-Native (BN)-SDS-PAGE analysis of thylakoid membrane preparations from *M. aeruginosa* PCC 7806 WT.

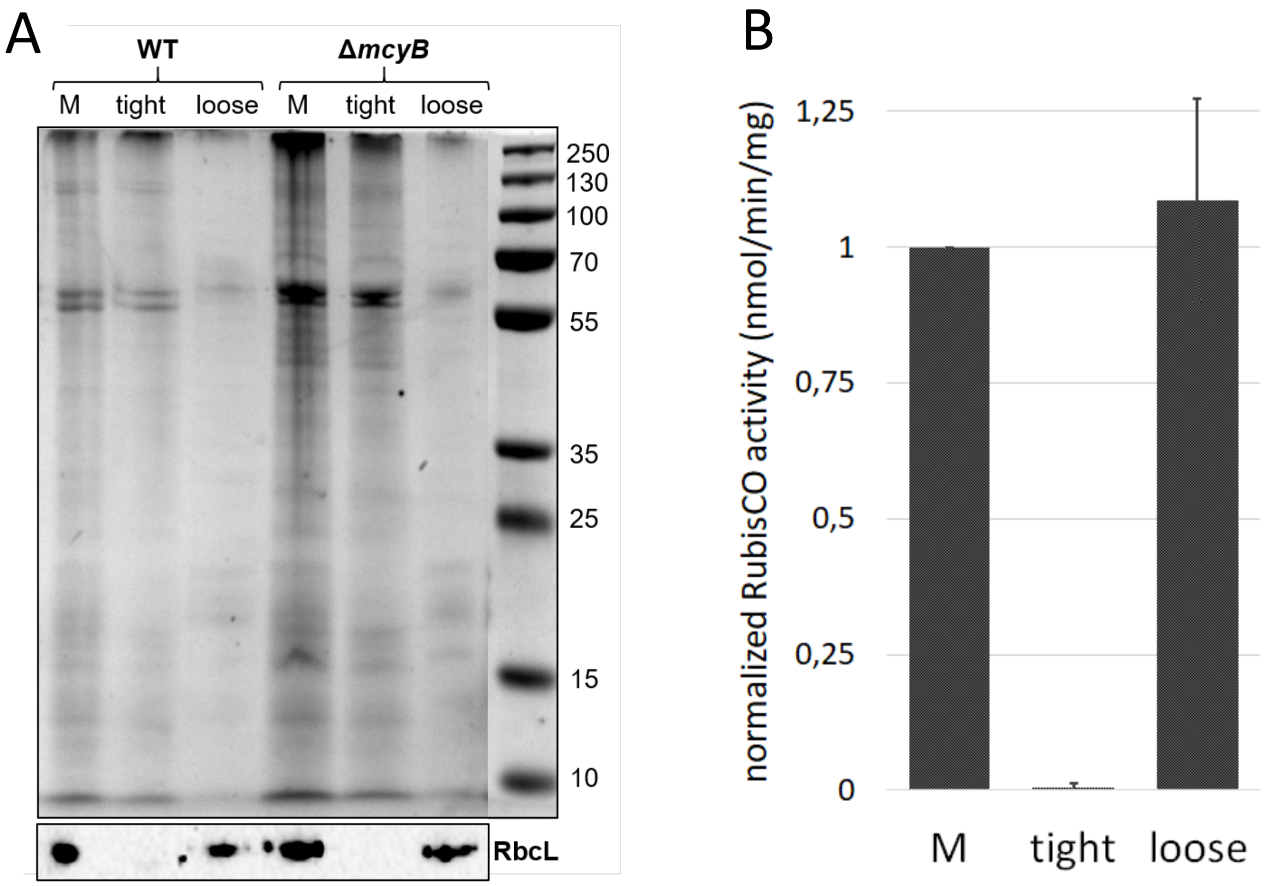


**Fig. S1** RbcL is loosely attached to membranes in *M. aeruginosa* PCC7806 and the ΔMcyB mutant. **A:** RbcL present in the membrane fraction (M) can be completely detached from the membranes by sonication (loose fractions). The remaining membrane fractions contain proteins more tightly connected (tight fractions) but lack detectable RbcL protein. **B:** RubisCO activity can only be detected in the whole membrane fraction (M) and in the loosely attached protein fraction. RubisCO activity was determined as nmol of fixed $C_i$ per minute and mg Rubisco and then normalized to the values of the M fraction.

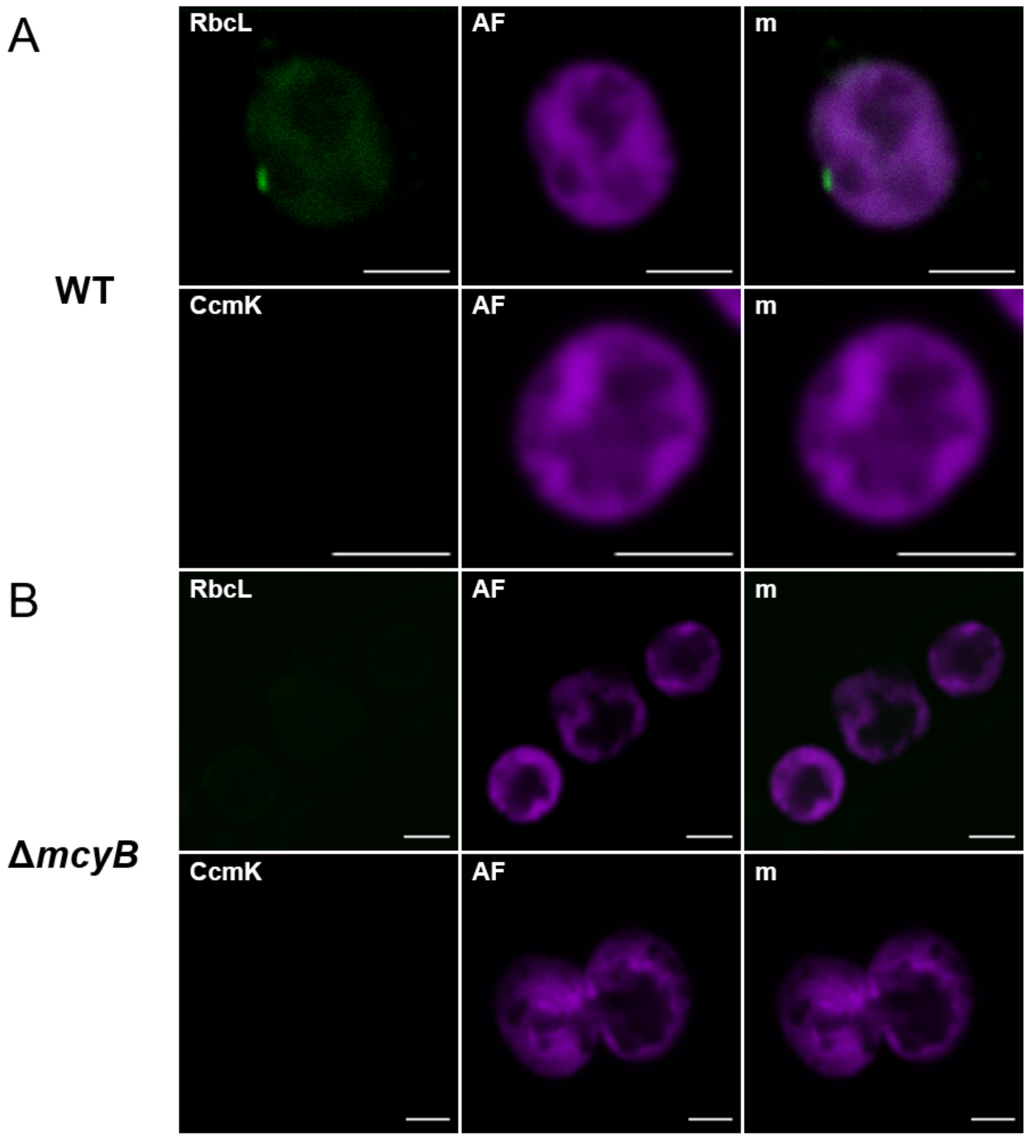


**Fig. S2** Immunofluorescence micrographs (IFM) showing the controls of the used secondary antibodies in *M. aeruginosa* wild type (WT) and Δ*mcyB* mutant IFM studies. **A:** Controls for *M. aeruginosa* WT show no specific signals for both proteins. Alexa Fluor 568 anti-Chicken was used as the secondary antibody for the visualization of RbcL. Alexa Fluor 488 anti-Rabbit was used as the secondary antibody for the visualization of CcmK. AF: red phycobilisome autofluorescence, m: merged image. **B:** Controls for *M. aeruginosa* Δ*mcyB* mutant show no specific signals for both proteins. Alexa Fluor 568 anti-Chicken was used as the secondary antibody for the visualization of RbcL. Alexa Fluor 488 anti-Rabbit was used as the secondary antibody for the visualization of CcmK. AF: red phycobilisome autofluorescence, m: merged image. Scale bar in all images: 2µm.

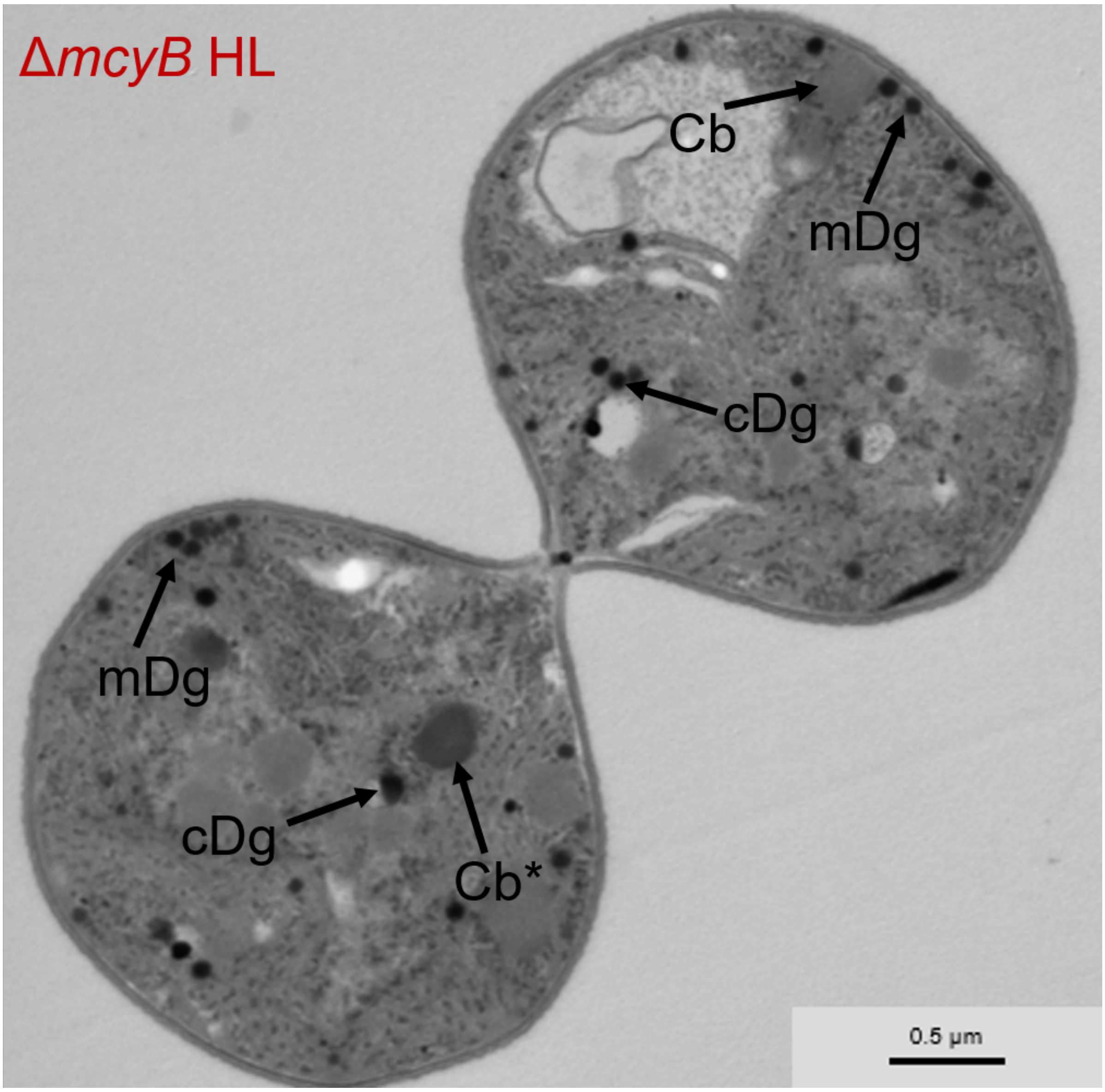


**Fig. S3** Transmission electron micrograph of *M. aeruginosa* PCC7806/Δ*mcyB* mutant grown towards high cell density ($OD_{750}$: 0.6) indicating presence of carboxysome structures with different electron densities. A carboxysome structure in the cytosol (Cb*) shows a higher electron-density than neighboring carboxysome structures. cDg – cytosolic dense granules likely representing RubisCO structures, mDg – membrane-associated dense granules likely representing RubisCO structures. Cb - carboxysomes

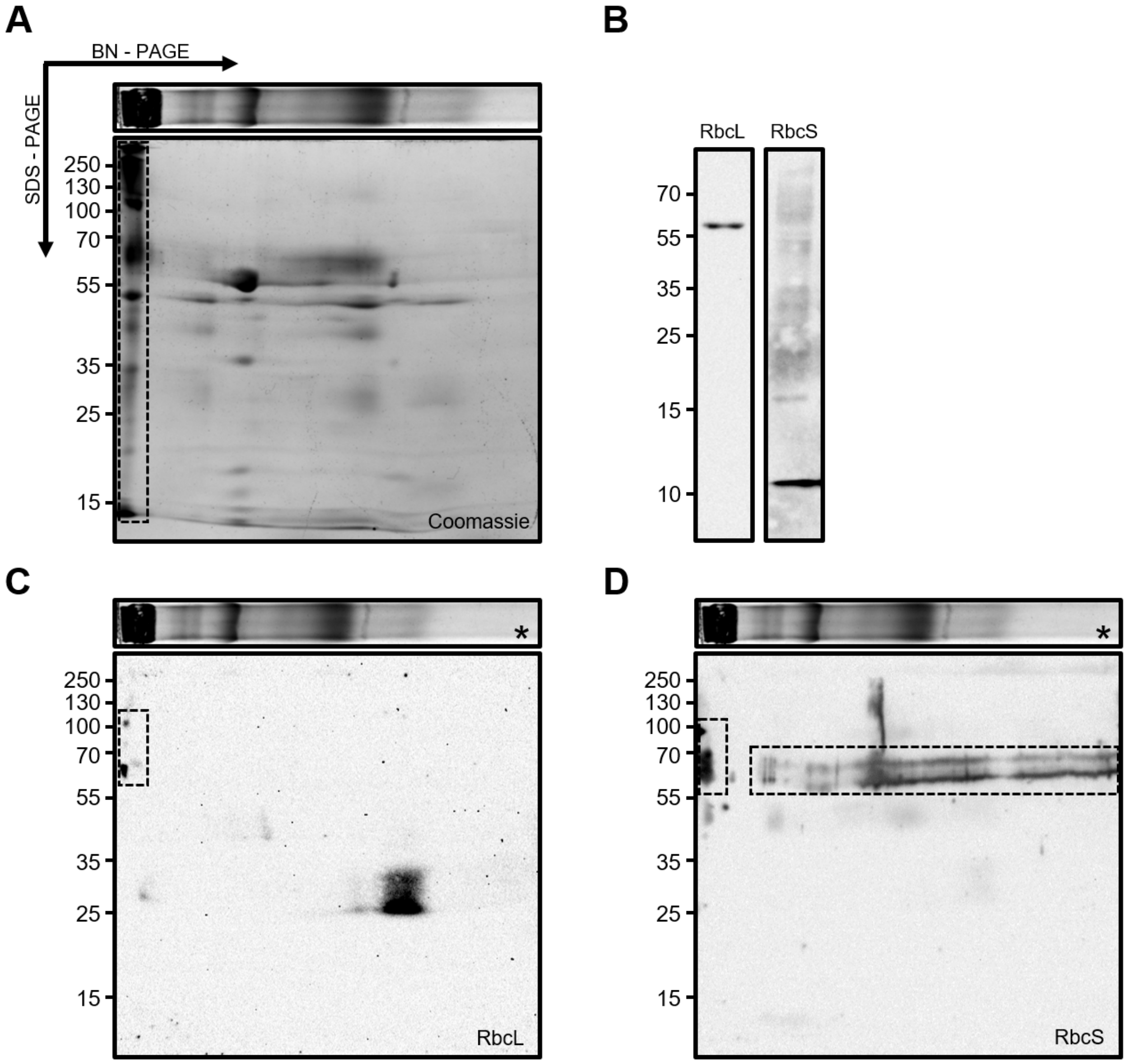


**Fig. S4** Blue-Native (BN)-SDS-PAGE analysis of thylakoid membrane preparations from *M. aeruginosa* PCC 7806 WT. A) Proteins were separated by BN-PAGE and SDS-PAGE analysis in the first and second dimension, respectively. B) One-dimensional SDS-PAGE analysis of thylakoid membrane protein fractions treated with 4M urea reveals presence of monomeric RbcS along with lower amounts of RbcL. C) 2D-Western Blot with anti-RbcL antibody reveals presence of RbcL in a high-molecular weight complex along with degraded RbcL in smaller size fraction. D) 2D-Western Blot with anti-RbcS antibody reveals presence of oligomeric SDS-stable RbcS in high molecular weight complex (HMW) along with oligomeric RbcS in various size fractions suggesting that RbcS unspecifically sticks to the membrane beside of its involvement in the HMW complex. The HMW complex comprising both RbcL and RbcS and RbcL-free RbcS fractions are highlighted with frames. The Coomassie-stained copy of the first dimension gel shown in Fig. 2A that was run in parallel with the BN gel stripes used for Figs. 2C and D is shown for orientation and indicated with an asterisk. Note that thylakoid membrane preparations may also contain proteins associated with the cytoplasmic membrane.
